# Accurate contact-based modelling of repeat proteins predicts the structure of Curlin and SPW repeats

**DOI:** 10.1101/809483

**Authors:** Claudio Bassot, Arne Elofsson

## Abstract

Repeat proteins are an abundant class in eukaryotic proteomes. They are involved in many eukaryotic specific functions, including signalling. For many of these families, the structure is not known. Recently, it has been shown that the structure of many protein families can be predicted by using contact predictions from direct coupling analysis and deep learning. However, their unique sequence features present in repeat proteins is a challenge for contact predictions DCA-methods. Here, we show that using the deep learning-based PconsC4 is more effective for predicting both intra and interunit contacts among a comprehensive set of repeat proteins. In a benchmark dataset of 819 repeat proteins about one third can be correctly modelled and among 51 PFAM families lacking a protein structure, we produce models of five families with estimated high accuracy.

**Author Summary:** Repeat proteins are widespread among organisms and particularly abundant in eukaryotic proteomes. Their primary sequence present repetition in the amino acid sequences that origin structures with repeated folds/domains. Although the repeated units are easy to be recognized in primary sequence, often structure information are missing. Here we used contact prediction for predicting the structure of repeats protein directly from their primary sequences. We benchmark our method on a dataset comprehensive of all the known repeated structures. We evaluate the contact predictions and the obtained models set for different classes of proteins and different lengths of the target, and we benchmark the quality assessment of the models on repeats proteins. Finally, we applied the methods on the repeat PFAM families missing of resolved structures, five of them modelled with high accuracy.

## Introduction

Repeat proteins contain periodic units in the primary sequence that are likely the result of duplication event at the genetic level [1]. Repeat proteins emerge through replication slippage [2] and double-strand break repair [3]. This protein class is present in all genomes but is more frequent in eukaryotic organisms [4–6] where they are involved in a wide range of functions [7]. In particular, due to their extended structures repeat proteins often behave as molecular scaffolds in protein signalling or for protein complexes as WD40 domain [8], or ankyrin repeats [9,10].

Repeat proteins are often conserved among orthologs [4,11] while exhibiting a more accelerated evolution and divergence among paralogs [11].

A classification of repeat proteins was proposed by Kajava [12,13] based on the length of the repeat units and the tertiary structure of the repeat units. According to Kajava’s classification, there are five classes of repeat proteins. However, in this study, we ignore class I and II because there are no available structures for class I, and class II structures are folded in a coiled-coil structure easy to be predicted. Moreover, the extreme amino acid compositional bias of many of these proteins makes it very hard to find the coevolving residues in these classes.

The dataset used in our study contains three classes of proteins divided into 20 subclasses divided by their secondary structure, according to RepeatsDB [14] Fig 1. The three classes are class III containing extended repeats (e.g. α and β solenoids), class IV containing closed repeats structures (e.g. TIM and β barrels and β-propeller), class V where the units appear as separate domains on a string. The units are also longer in class V than in other classes.

**Figure 1.**
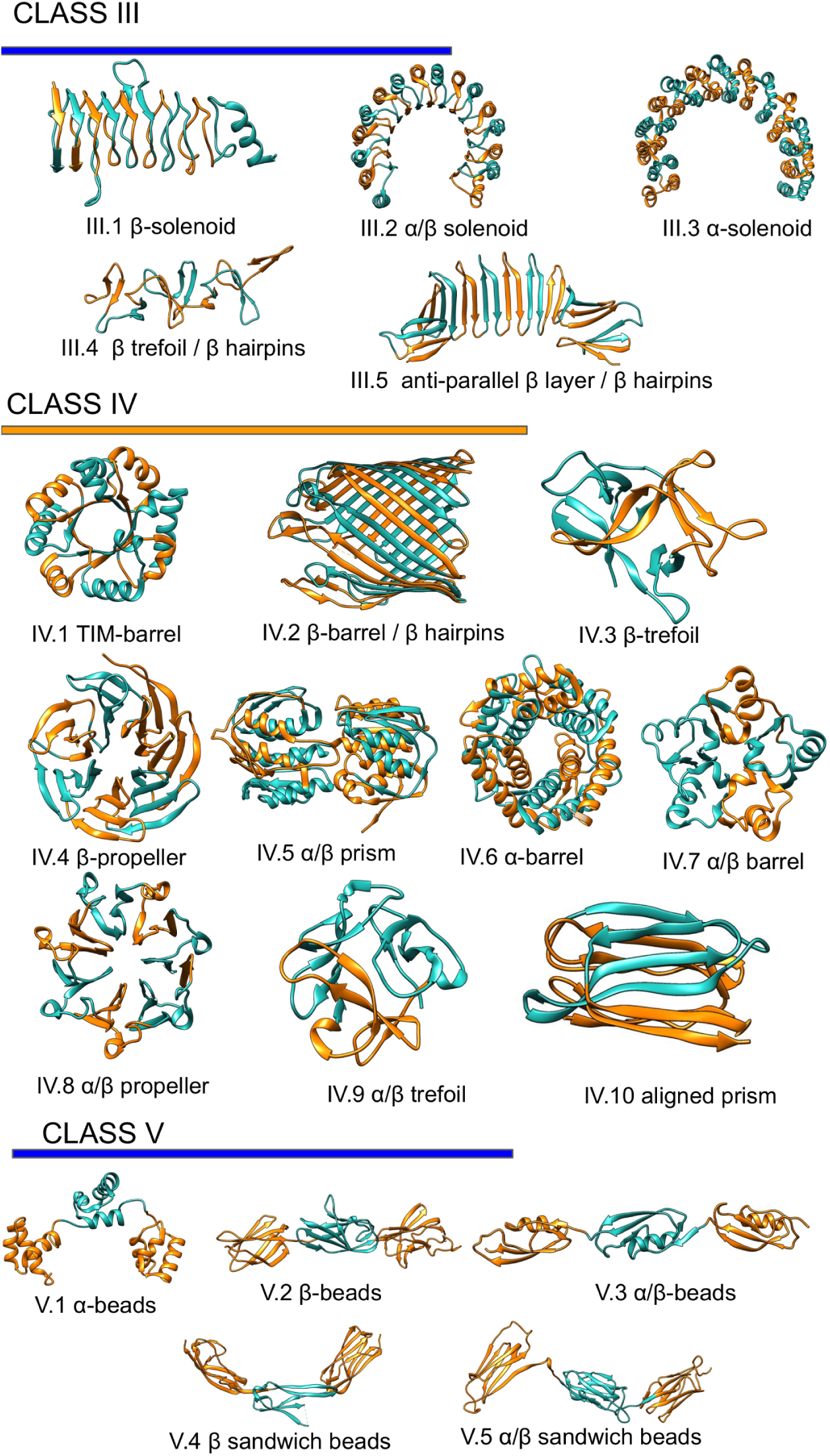
Repeats proteins classification. Representation of the repeats classes and subclasses as classified in repeatsDB 2.0 [14]

Class III is dominated by solenoid structures (Figure1 III.1, III.2 III.3) [13], and there is a wide range in the numbers of units (from 4 to 38). Also, the length of the individual unit is widely variable, e.g. β-solenoid have significantly shorter repeats compared with α and α/β solenoid [13]. Two subclasses: β-trefoil/β-hairpins, anti-parallel and β-layer/β-hairpins form extended beta strands without the bend typical of the solenoid.

Members of class IV are constrained in variability by the closed fold. Indeed despite ten subclasses of different units fold the number of units go from 3 to 16, and the proteins with more than ten units are rare. The length of the units is in between class III and V [13].

Class V has the longest units, which fold into proper domains and also a low number of units with few interactions between them.

However, many repeat proteins lack a resolved structure or a template to perform homology modelling. Residue-residue contact prediction is the most promising template free method [15]. Contact prediction methods identify residues co-evolution from multiple sequence alignment and identify the evolutionary constraints of the residues imposed by the tertiary protein structure [16]. Nevertheless, repeat proteins are a difficult target for contact prediction; the internal symmetry introduces artefacts in the contact map at a distance corresponding to the repeated units [17].

Here, we benchmark the deep-learning-based contacts prediction program PconsC4 [18] against the GaussDCA [19] on a comprehensive dataset generated from RepeatsDB [14]. The predicted contacts were then used as constraints to generate proteins model, and their quality was then tested by Pcons [20]. On the base of the benchmark, we propose models for the protein structures of PFAM protein families missing of resolved structures.

## Results and Discussion

### General contact prediction analysis in repeat proteins

To assess the quality of the contacts predictions among repeat protein classes, we generate a dataset of proteins, clustering at 40% of identity, the reviewed entries of RepeatsDB [14]. For each repeats region present in the dataset we extract the sequence of a representative repeat unit and a pair of repeats, obtaining in this way three datasets: i) a single unit datasets; ii) a double unit datasets; iii) complete region datasets.

For all the three sets of proteins, multiple sequence alignments (MSA) and secondary structure predictions were generated. Subsequently using the MSA as input for PconsC4 and GaussDCA [19] contacts were predicted for each family. The performance of the contact predictions was evaluated for each subclass separately. As expected, PconsC4 over-perform GaussDCA in all the three sets and all the classes of repeat proteins, Figure 2.

**Figure 2.**
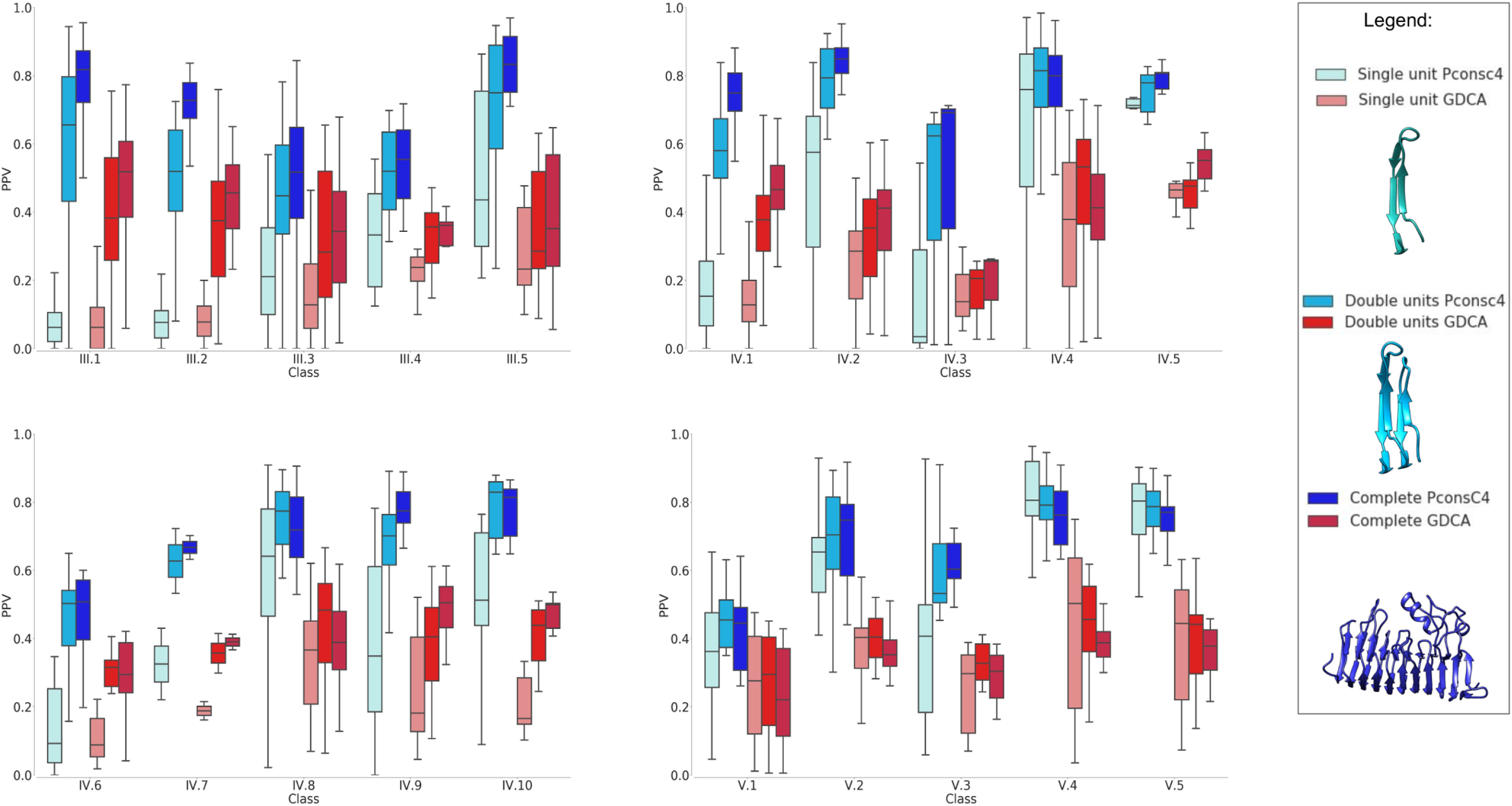
Precision of contact predictions. Positive Predicted Value (PPV) for the GaussDCA (red) and Pconsc4 (Blue) contact prediction for each subclass. In light colour the single unit dataset, in medium colour the double units dataset, and in dark colour the complete region dataset.

Here, it should be remembered that PconsC4 use the GaussDCA prediction as an input for the U-net [32] that learn to recognize specific contacts patterns [18].

In general, the predictions for the full length regions (darker colors in Fig. 2) give better results than split the proteins into units but with some exceptions. In particular in class V, that is composed by bigger units forming repeats of the *“beds on a string*” type, the splitting in units may help, especially in some subclasses, to reach better contacts prediction performance as discussed later.

Furthermore, PconsC4 appear efficient in removing the DCA repeats artefacts compared with GaussDCA. In Fig. 3 are shown some contact maps examples. In the GaussDCA predictions are evident the periodic artefacts of wrong predictions (red dots) forming perpendicular lines. These appear to be contacts between equivalent positions in the repeat unit.

**Figure 3.**
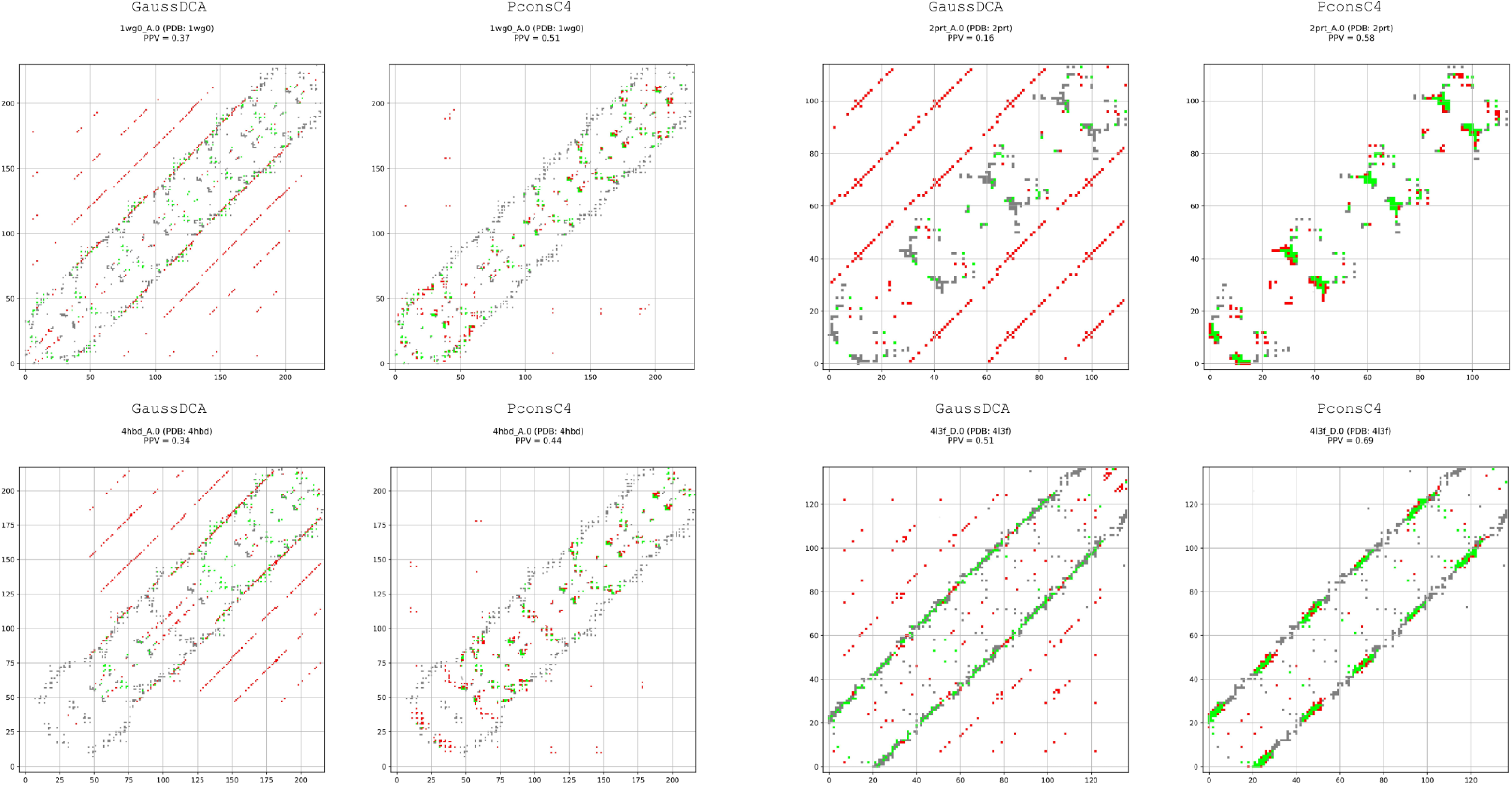
GaussDCA and PconsC4 contact maps. Contact map for a prediction with a) GaussDCA b) PconsC4. In grey, the real contacts from the structure, in green the corrected predicted value, in red the false predicted value.

Finally, it is well known that the quality of the prediction is directly correlated with the number of sequences in the starting MSA, especially for the DCA methods [18]. The same trend is confirmed among protein repeats, Fig. 4, where the repeats with a smaller MSA are predicted with lower PPV. PconsC4 and GaussDCA show the same pattern in the average PPV except for an increase of the PPV for PconsC4 with MSA with a Neff Higher than 12.

**Figure 4.**
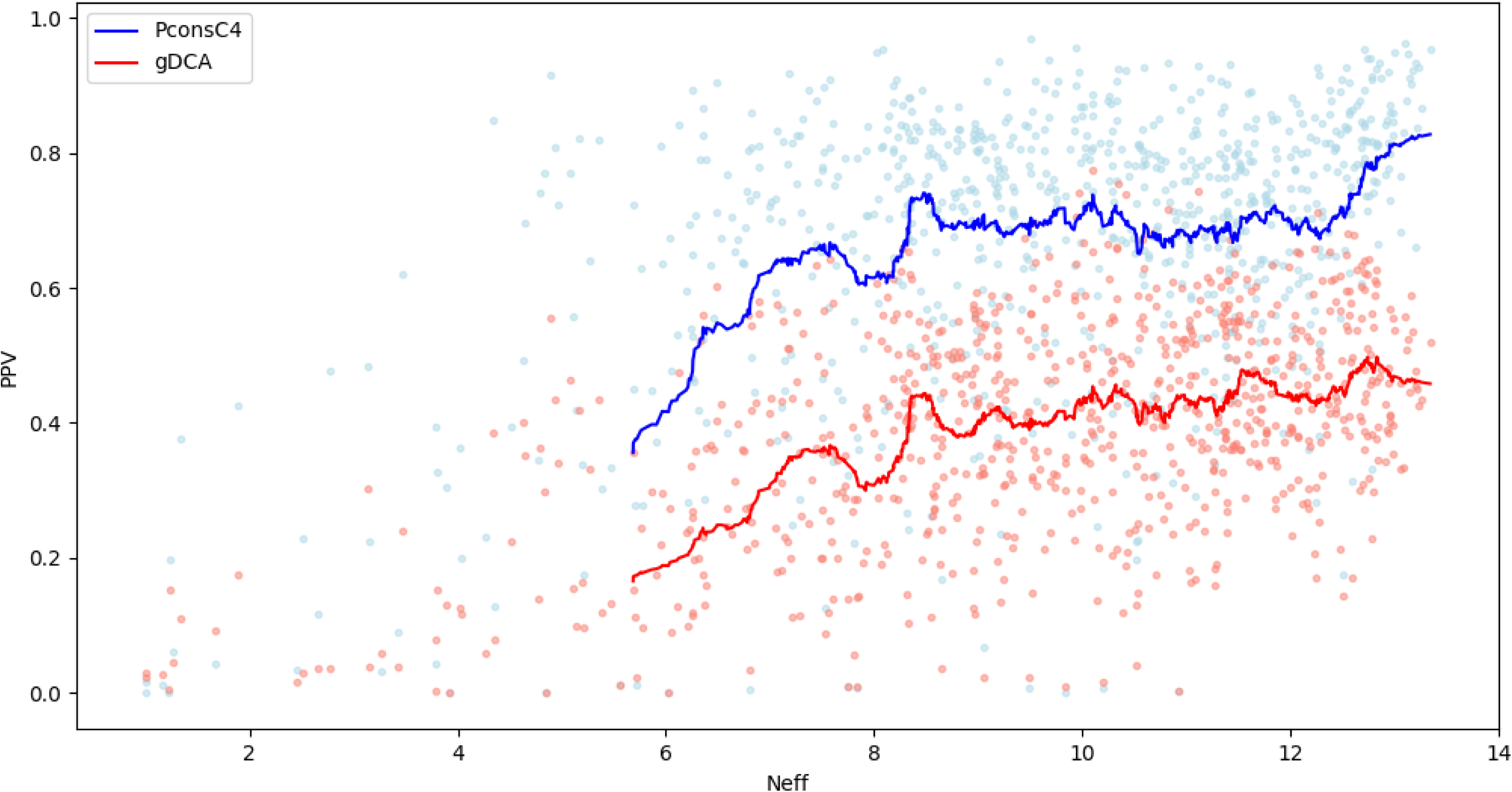
PPV versus Neff. Positively Predicted Value for GaussDCA in red and PconsC4 in Blue on the Neff value (number of effective sequences length weighted with length).

### Differences among repeat classes in contacts prediction

Fig. 2 shows variations in the percentage of correct contacts among different protein repeat classes and subclasses, to clarify the origin of these differences, we investigated more in-depth the origin of the predicted contacts.

One central aspect that affects the difficulty of prediction is due to the pattern of contacts [33]. In general, contacts that are parts of larger interaction areas are better predicted as well as interactions between residues that are close in the sequence. A comparison between the intra-unit and inter-unit contacts are shown in Fig 5a. Here, we obtained the number of intra and inter-unit contacts from the PDB structures and we selected the same number of intra and inter units from the contact predictions. The PPV was finally calculated as the number of correct contacts over the number of the selected contacts.

**Figure 5.**
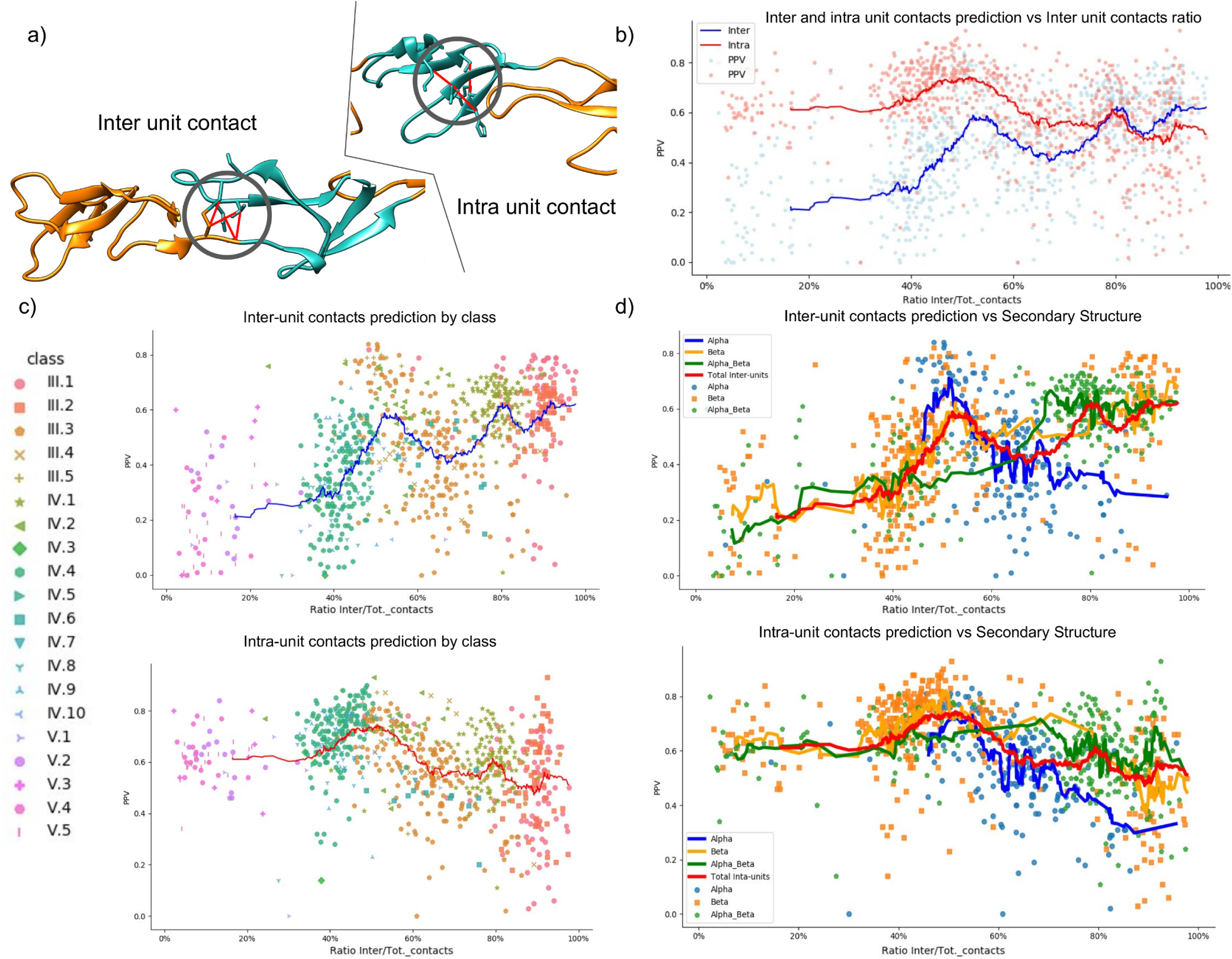
Predicted contacts analysis. Positive Predicted Value (PPV) obtained by PconsC4 for different types of contacts. a) Examples of inter- and intra- unit contacts. b) In red, the PPV for intra-units contacts in blue PPV for inter-units contact. c) Repeats subclasses. In red, the PPV for intra-units contacts in blue PPV for inter-units contact, colors and shapes in the scatter plot indicate different protein subclasses. d) Secondary structure. In red, the overall PPV, in blue, the α-helical subclasses, in green, the α-helix/β-strandβ-strand subclasses, and in orange, the β-strand subclasses.

On average the intra-units contacts are predicted with higher accuracy than the inter-unit one, but this is not true for all protein classes. This behaviour is due to significant differences of the units structures among the classes: in class III the unit are short, and the residues form contacts mostly with the neighbour units; in class V, on the contrary, the units are long, folded in independent domains and the contacts are predominantly inside the units with few inter-unit contacts; class IV is halfway between class III and V. The inter units contacts of class III and partially of class IV results easier do be predicted then class V ones, because they form clearer patterns in contact maps. On the contrary, the intra-unit contacts of class V are predicted better than class III and IV for the same reason. We plot the PPV versus the ratio of the inter-unit contacts over the total number of contacts of each protein. The PPV show an inverse relation with the ratio of inter-unit contact of the protein (Figure 5.b,c,d).

The inter-units PPV is low for the proteins with an inter-contact ratio lower than 20% constituting class V. Figure 5c shows the lowest inter-units contacts PPV while the PPV between inter- and intra-contacts invert the trends at a ratio of 80%. This switch corresponds to solenoid structures and TIM Barrel that have a ratio between 80%-100% larger interaction surfaces between different units than inside a single unit.

In Figure 5d, we divided the proteins into their secondary structure class. Proteins subclasses containing only β-strand or α-helix/β-strand appear easier to predict. The plots show a steep decrease in the PPV values around the ratio of 50% helix for both intra- and inter-units contacts. This is due to the α-helix subclasses component. α-helix are harder to predict because they produce a less clear contact pattern compared with β-strand.

### Protein model generation and quality assessment

Proteins models were generated using CONFOLD [28] starting from the contact predictions of PconsC4 and the PSIpred secondary structure as constraints. In Fig. 6 we compare the TM-score between the first model ranked by CONFOLD and the corresponding PDB protein structure.

**Figure 6.**
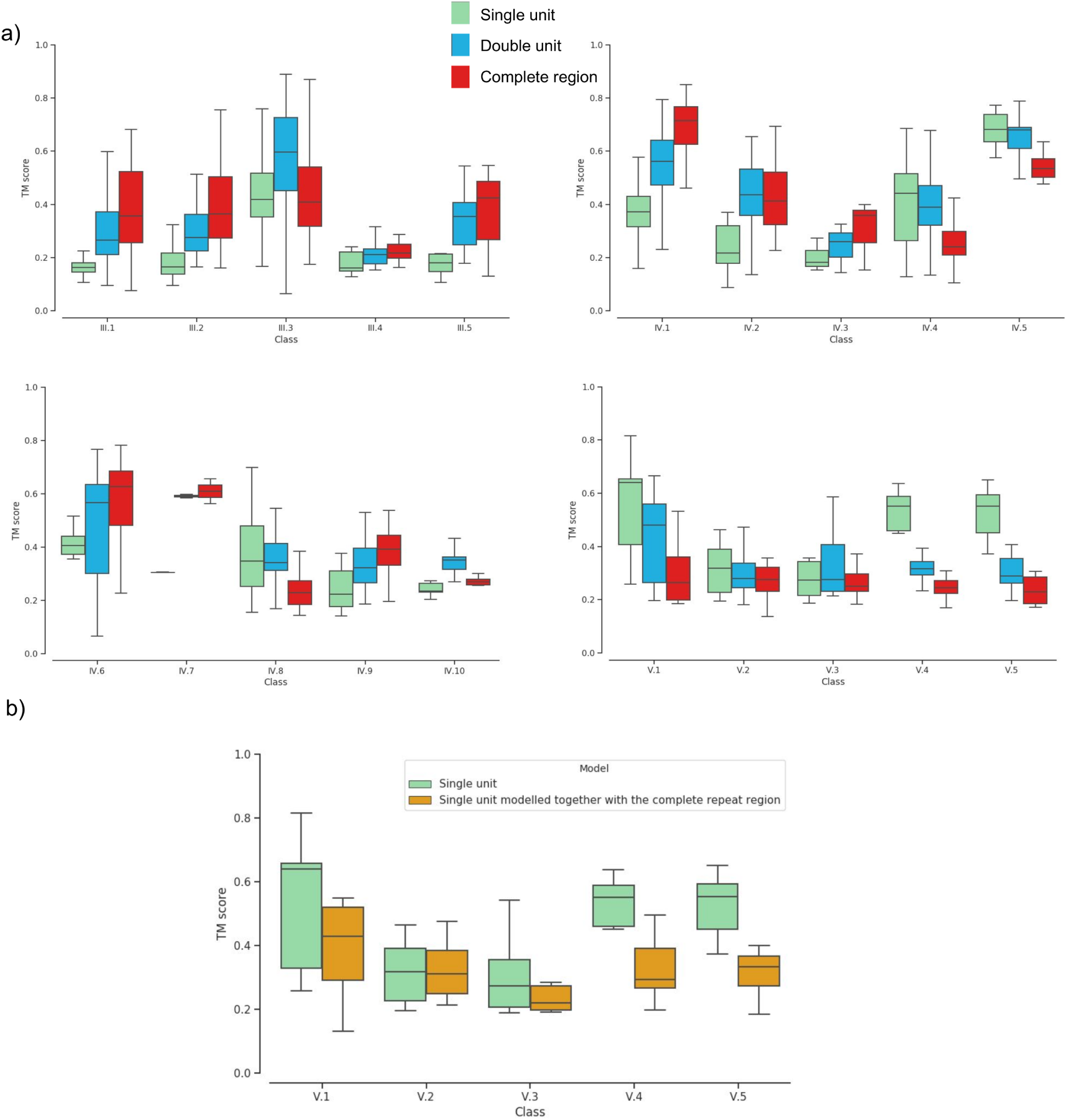
Protein model quality. a) TM-score in all the subfamilies. In sea-green the single unit prediction, in blue the double units prediction, in red the complete region prediction. b) TM-score of the subfamilies of class V. In green the single unit prediction and in brown the prediction of that unit when the entire region is modelled.

Although the best contact predictions were, on average, obtained with the complete regions, still splitting the structure lead in some cases to a better model; this is true in particular for the “Beads on a string” class V, but single-unit models are also useful in the “propeller” subclasses class IV: IV.4 β propeller, IV.8 α/β propeller and IV.5 α/β prism. Moreover modelling a couple of units lead to the best models in two subclass III.3 α solenoid IV.10 and aligned prism. All these subclasses except α-solenoid have a low ratio of inter-units contacts (below 50%) Fig. 5b, however α-solenoid where the complete protein reaches in some cases a length of 1000 residues. Moreover, the bend of the protein is very difficult to predict, and the models result in a series of straight helices.

It is questionable if the lower quality of the models of the complete region is due to a general decrease in the performance or only to the impossibility to model the correct interaction among different domains. To answer this question, we analysed more in deeper class V, where the decrease in the performance is most evident. We extract from the “complete region model” the same units and the units previously modelled as single and double units Fig. 6b. Interestingly, even the single units and the double units extracted from the complete region modelling have a lower or similar accuracy compared with the single and double units modelled alone, Figure 6b. This is observed is regardless of the quality of the prediction of the contacts in the complete region prediction, Fig. 2, suggesting that the poor performance is not only due to the more difficult prediction of the interdomain contacts but also due to a limitation of the modelling of longer proteins.

In order to evaluate the model quality, we plot the TM-scores of the models against the score obtained from the quality assessment method Pcons [20], Fig. 7. In light of this result, we consider the models of a complete repeats region reasonably correct when they reach a Pcons score of 0.4. The complete dataset with Pcons prediction is reported in Table S1.

**Figure 7.**
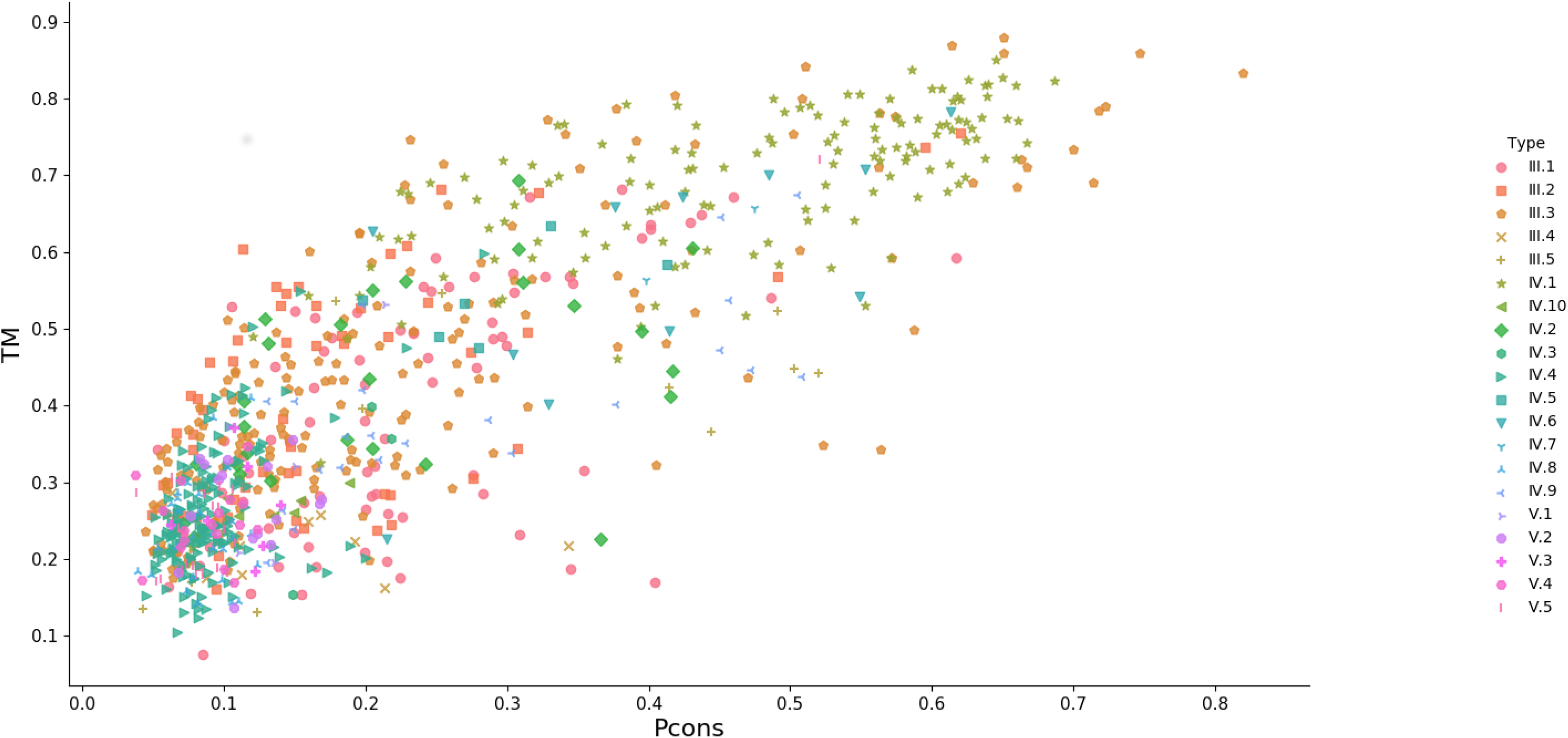
TM-score versus Pcons-score. TM-score versus Pcons-score for complete region models.

### Modelling of repeat proteins without resolved structures

In order to predict the structure of new repeats families, we selected 51 PFAM repeats family without resolved structure. A representative sequence of each family was run against Uniclust30 with HHpred, and the resulting MSA was used to predict the contact map that was used together with the PSIpred prediction as constraints to generate the models.

All the models were evaluated with Pcons, but only five of them reach a Pcons score higher than 0.4. These are the PFAM family; MORN 2, SPW, Curlin rpt, RTTN N, RHS repeat, Table 1 (In Supplementary the target/template alignments).

In order to further prove the reliability of these models and perform a more comprehensive protein modelling approach, we associated homology modelling and the contact-based modelling approach. For three out of five proteins, HHsearch returned a highly reliable template, Table 1.

| PFAM Family | Rappresentative sequence (Uniprot ID) | Pcons score | Template with seq. coverage > 70% (PDB ID) | HHsearch probability | TM score contact model/homology model |
| --- | --- | --- | --- | --- | --- |
| **MORN 2 (PF07661)** | **Q8RH85** | **0.711** | **1MUF_A** | **99.37** | **0.5003** |
| SPW | A0A2A3HD64 | 0.674 | 5EQC_A | 29.35 | / |
| Curlin rpt | Q8EIH3 | 0.576 | 2N59_A | 1.97 | / |
| **RTTN N (PF14726)** | **W5P499** | **0.490** | **4U2X_E** | **94.03** | **0.3529** |
| **RHS repeat** | **A0A1G0MXS8** | **0.407** | **5KIS_B** | **99.46** | **0.5823** |

In Fig. 8, the superimposition between homology modelling and contact based model is shown. In all three the protein family there is a substantial agreement between the two approaches. MORN 2 family contact-based and homology model are in agreement except for loops and the bend of the central beta-strand.

**Figure 8.**
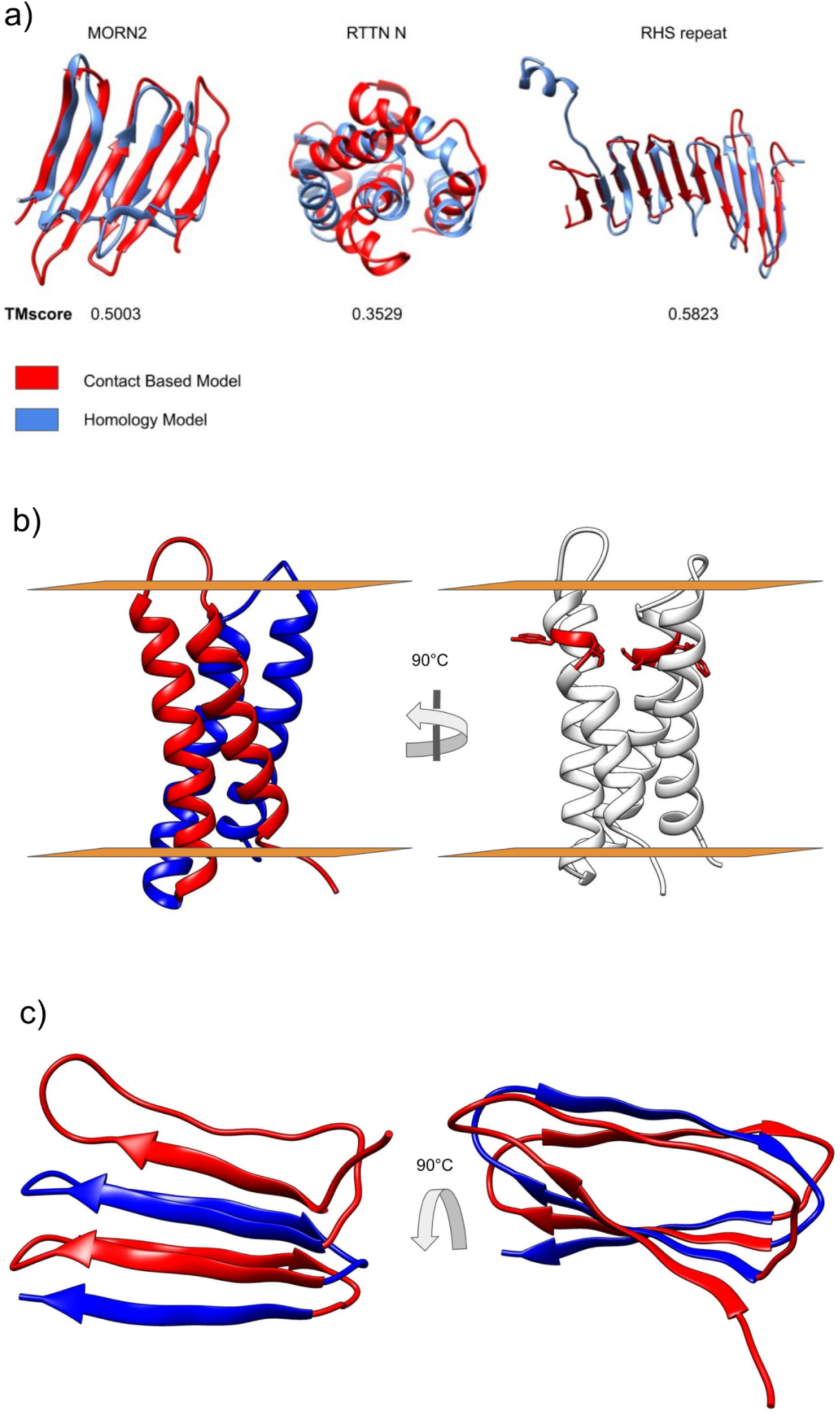
High quality protein models. a) Superimposition between the contact-based models and the Homology Model performed with Chimera [34] and their respective TM- score. In red, the contact-based models and in light blue Homology models. b) Protein model of SPW family in the membrane (light brown). On the left in blue and red the two repeated units on the right in red the SPW motif. c) Protein model of Curlin repeats, in blue and red the repeated units.

The RTTN N is the family showing the lowest TM-score between the two models mostly due to a different rearrangement of the firsts three alpha-helices. Has to be mentioned, however, that despite a high probability score, the identity between the target and the best template is only 7% (Figure S1b) making hard to determine which is the best model.

In RHS repeat family, the score between the contact-based models and the homology model share a TM-score of 0.58. Only the N terminal is modelled in a different with an extra beta-strand in the contact-based model and an alpha helix in the template-based modelling. However, we argue that in this case, the contact-based modelling overperform the Homology model; indeed the contact prediction mode is in agreement with the secondary structure prediction that predicts an N-terminal Beta strand (Figure S1c).

The remaining two PFAM families do not have suitable templates, and contact-based modelling is the best suitable method for model them.

### SPW family

According to the PFAM database, the SPW family is present in Bacteria and Archaea in one or two units, and in a few cases in association with a Vitamin K epoxide reductase or NAD-dependent epimerase/dehydratase domain. Each repeated unit is formed by two transmembrane alpha-helices and is characterized by an SPW motive [35]. According to our model, the repeated motifs is buried in the membrane symmetrically located close to the extracellular side, Fig. *8b*. PFAM architectures show many proteins with only a single SPW motif however a more careful analysis of these sequences shows that in many cases they contain a second degenerate SPW unit before or after the one identified where however the proline residue is conserved (Figure S2).

The Tryptophan is on the outer side of the protein facing the bilayer while the proline is on the inner side of the protein promoting the formation of a kink in the transmembrane helix [36]. The motif “SP” in particular, increase the bending effect of proline significantly due to their hydrogen bond pattern [37], indeed due to the structural propriety, the motif is relatively rare in membrane proteins [37].

### Curlin repeats family

Our model results in a β-solenoid structure, Fig. *8c* DeBenedictis et al. in 2017 presented and discussed ab initio models for the Curlin repeats family members CsgA and CsgB [38], their best models is in agreement with our model (a direct comparison is difficult as the coordinates is not available of their model). The model is furthermore confirmed by the partial structure of the repeat units of CsgA published by Perov et al. [39] where they crystallize in parallel β-sheets with individual units situated perpendicular to the fibril axis (corresponding PDB IDs are 6G8C, 6G8D, 6G8E).

## Conclusion

The modelling of the unknown PFAM families was challenging. Only 10% of the datasets had a Pcons score equal or higher to 0.4; compared to 21% in the benchmark dataset. However, the differences between the two datasets have to be taken into account. It is known that a smaller MSA affects the prediction of contacts and known structures are biased towards the larger family [40] Indeed our “Unknown protein families” dataset shows a significant lower Neff score compared with the PDB benchmark set, Figure 9a. Moreover, in the Unknown protein set, there are more eukaryotic-specific protein families (Fig. 9b).

**Figure9.**
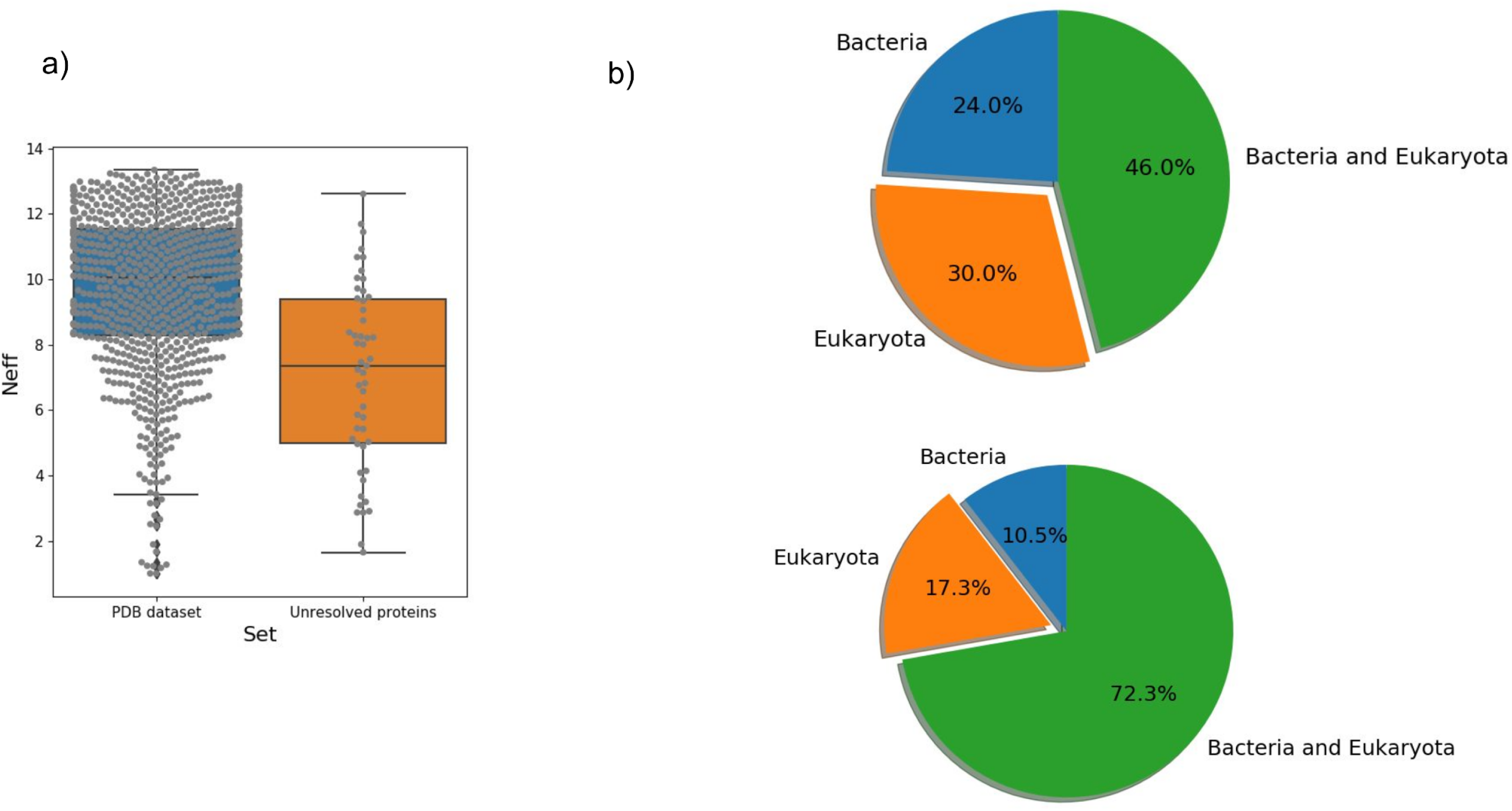
Datasets comparisons. a) Neff score comparison between the two datasets. b) The variation in the membership to the three domains of life between the PFAM families of the “Unresolved Proteins Dataset” and the “PDB dataset”.

Despite the significant improvement brought by deep-learning in contact prediction, there is still room for improvement. The prediction of inter-domain contacts accuracy is often lower than the intra-units one and the development of a model trained explicitly on repeats protein datasets might improve the result. Furthermore, the folding part of the pipeline is a limiting step, in particular for long proteins.

In our study, we performed a comprehensive coevolution analysis on repeat protein families, and we show that PconsC4 contact-predictions method overcomes the traditional difficulties of DCA methods for this class of proteins. We investigated the modelling of repeat units, and we provided a “titration curve” for Pcons score for repeat proteins. Finally, we test our pipeline on PFAM families without protein structures showing its usefulness in providing new structural information.

## Materials and Methods

### Datasets generation

The repeat protein dataset was generated starting from the 3585 reviewed entries in RepeatsDB [14,21], http://protein.bio.unipd.it/repeatsdb-lite/dataset. The proteins of class I and II were removed, and then the dataset was homology reduced using CD-HIT [22] at 40% identity resulting in 819 repeats regions. From this “complete region dataset” two others datasets were generated: I) A “single unit” dataset with one repeat unit for each region; II) A “double unit” dataset with a pair of units per each repeat region. In the two derived datasets, the representative units were selected, avoiding or at least minimizing, the presence of insertions.

The non-resolved repeats protein family dataset was generated, collecting all the repeat proteins families with missing structural information present in PFAM [23] in May 2019 and removing the domains with a significant overlap with the disorder prediction. It results in 51 protein families. The representative sequence for each family of repeat was chosen for matching these criteria: 1) select the most common architecture; 2) Include when possible at least three repeat units.

### Multiple sequence alignment (MSA)

The multiple sequence alignments (MSA) were carried out using HHblits [24] with an E-value cutoff of 0.001 against the Uniclust30_2017_04 database [25]. The number of effective sequences of the alignment, expressed as Neff-score, was calculated by HHblits and used for subsequent analysis.

### Contact prediction and models generation

The protein models were generated following the PconsFold2 protocol of [26]. The secondary structure of the repeat regions was predicted by PSIpred [27]. Protein contacts were calculated with PconsC4 [18] and together with the secondary structure predictions were used as input for Confold [28]. The modelling was run using the top scoring 1.5 L contracts where L is the length of the modelled regions and the two-stage modelling.

### Contacts analysis

A protein contact was defined as two residues having a beta carbon distance equal or lower than 8Å in the PDB structure and farther than 5 residues in the sequence. Using this definition, we assess the number of correctly predicted contacts (the Positively Predicted value (PPV)) taking into account the top-scoring 1.5 L contracts.

In the intra/inter unit contacts analysis, the predicted contacts of each protein were divided between i) intra-unit contacts, if between residues inside the same unit; ii) inter-units if the residues are in different repeat units. The units mapping was taken from the RepeatsDB database [14]. In this analysis, we calculate the number of intra- and inter-unit contacts existing in the PDB structure, and we selected the same number of intra- and inter-units predictions. The PPV was then calculated as the number of correct contacts over the number of the selected contacts.

### Homology modelling

Templates for homology modelling were searched by HHsearch [29] using the HHpred web-server with default settings on PDB_mmCIF70_3_Aug database. Subsequently, the models were generated by HHpred [30].

### Protein models analysis

The model quality was assessed using Pcons [20]. We download and installed Pcons. With the option -d we predicted the quality among the model in the stage2 folder generated by Confold. Pcons uses a clustering method, and the score is simply the average structural similarity to all models, as measured by the S-score.

The TM-score was calculated using TMalign [31]. To ensure that the protein structure and the model were properly aligned the option -I was used, providing a local protein alignment for the two sequences.

## Supporting Information

**Table S1.** Unknown protein family dataset. In the columns are reported respectively: the UniProt ID of the modelled sequence, the PFAM family, the Pcons score.

| Uniprot entry | Pfam | Pcons_score |
| --- | --- | --- |
| S6TLB9 | PF14882 | 0.071 |
| W5U916 | PF15907 | 0.072 |
| D7MCA5 | PF07725 | 0.079 |
| A0A2A2LSA2 | PF14625 | 0.08 |
| R5P8A5 | PF07538 | 0.081 |
| G1XIQ8 | PF13446 | 0.083 |
| A0A0B0HSH2 | PF13753 | 0.094 |
| J1S4N0 | PF11966 | 0.094 |
| U5QIU9 | PF06739 | 0.096 |
| A0A1A9WU23 | PF02363 | 0.102 |
| A0A252E8A5 | PF17660 | 0.103 |
| L8TNF3 | PF08310 | 0.108 |
| W4LGN0 | PF14312 | 0.111 |
| U7Q0S5 | PF10281 | 0.116 |
| D3UXB8 | PF07634 | 0.117 |
| Q29AL9 | PF14939 | 0.121 |
| A0A1Z4C3E9 | PF17164 | 0.122 |
| Q9NZT2 | PF04680 | 0.123 |
| D5SU36 | PF07639 | 0.139 |
| G3VIY2 | PF00400 | 0.143 |
| D3BR65 | PF00526 | 0.15 |
| I3BT02 | PF03640 | 0.153 |
| A0A094KVK3 | PF00880 | 0.159 |
| R6YH89 | PF14903 | 0.16 |
| O64827 | PF18868 | 0.164 |
| A0A257INW4 | PF13573 | 0.165 |
| A7S4G3 | PF07016 | 0.167 |
| R2SEH8 | PF18780 | 0.168 |
| T0NQR8 | PF08043 | 0.176 |
| A0A1V1NWB1 | PF08309 | 0.179 |
| A0A0L7M9B8 | PF07981 | 0.188 |
| A7RAI0 | PF06598 | 0.2 |
| G1RYA9 | PF06049 | 0.209 |
| S7J9T7 | PF02415 | 0.215 |
| V5CQL0 | PF03406 | 0.232 |
| A0A1I7SWM5 | PF00839 | 0.251 |
| Q7RTC2 | PF12135 | 0.252 |
| Q8PXT0 | PF06848 | 0.253 |
| W5N853 | PF03128 | 0.253 |
| Q6YQH3 | PF11178 | 0.262 |
| R7MCC4 | PF13475 | 0.278 |
| Q6P6X2 | PF10578 | 0.295 |
| H9GJM4 | PF13330 | 0.319 |
| U2FCE1 | PF12779 | 0.355 |
| F3GDU0 | PF00818 | 0.392 |
| A0A1G0MXS8 | PF05593 | 0.407 |
| W5P499 | PF14726 | 0.49 |
| Q8EIH3 | PF07012 | 0.576 |
| A0A2A3HD64 | PF03779 | 0.674 |
| Q8RH85 | PF07661 | 0.711 |
| W5Q8K9 | PF15390 | 0.079 |

**Figure S1.**
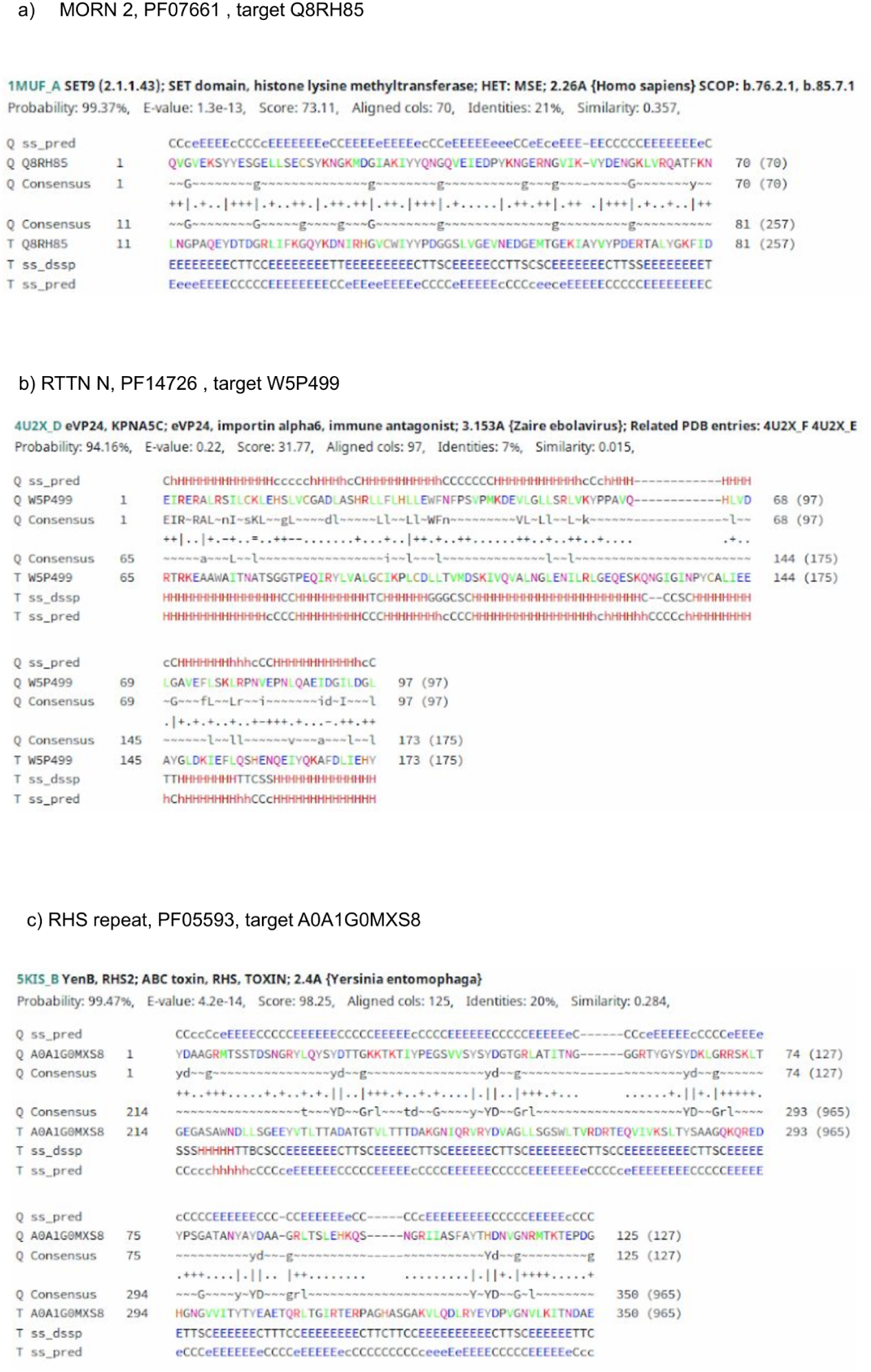
Target/template alignments. Target/template alignments for the homology modelling.

**Figure S2.**
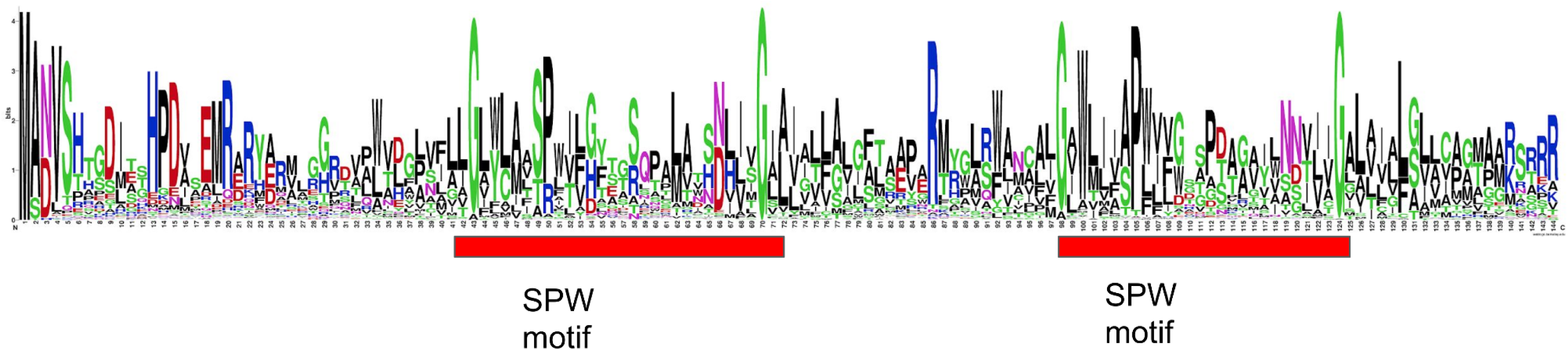
Amino Acid frequency of the single domain architecture sequences. From the logo is possible recognize two SPW domains, one of them degenerated (in particular the first Serine in the second motif) that is not recognized by PFAM.

